# Impacts of perinatal factors on white matter outcome at 8 to 10 years by diffusion tensor imaging

**DOI:** 10.1101/2025.10.05.680580

**Authors:** Injoong Kim, Omar Azrak, Mark Foster, Emil Cornea, Sang Kyoon Park, Yoonmi Hong, Martin Styner, John H. Gilmore

## Abstract

**Background:** While perinatal factors are known to influence brain development, their long-term impact on white matter microstructure remains incompletely understood. Previous studies using tract-based spatial statistics (TBSS) have shown limited associations between neonatal measures and later white matter development.

**Methods:** We investigated associations between perinatal factors (birth weight [BW], gestational age [GA], and head circumference [HC]) and white matter microstructure in 117 children aged 8-10 years from the UNC Early Brain Development Study cohort. Diffusion tensor imaging (DTI) data were analyzed using a fiber tract-based framework examining 54 major white matter tracts. Statistical analysis was performed using a functional analysis of fiber tract profiles.

**Results:** GA and BW showed widespread significant associations with white matter microstructure (38 and 36 out of 54 tracts, respectively), while HC showed limited associations (3 out of 54 tracts). Post-hoc univariate analysis revealed stronger associations with axial diffusivity (AD) compared to radial diffusivity (RD) or fractional anisotropy (FA). AD associations with BW, GA, and HC were found in 30, 31, and 8 tracts, respectively.

**Conclusions:** Using a fiber tract-based analysis approach, we demonstrated that GA and BW are strongly predictive of white matter organization at school age, while HC showed limited predictive power. The predominant associations with AD suggest these perinatal factors primarily influence axonal organization rather than myelination. These findings enhance our understanding of how early life factors impact long-term brain development.

## Introduction

Pre and postnatal brain development is a complex process that is under active investigation, and it is a continuous and dynamic process that extends well into early adulthood (1). In the last two decades, advances in magnetic resonance imaging (MRI) provided researchers with tools to investigate developmental changes of cerebral gray and white matter. Diffusion tensor imaging (DTI) has shown to capture white matter development that intrinsically linked with changes in the brain’s efficiency, which is related to a variety of cognitive, behavioral, and psychopathological outcomes (2-4), DTI reveals white matter microstructure by using the orientation and integrity of white matter tracts (5), enabling the examination of tissue microstructure in vivo, and offering quantifiable data that are relevant to both brain development and injury (6). Metrics obtained from the DTI are highly sensitive to developmental shifts and white matter changes associated with prematurity and perinatal risk factors (7-10). Building upon this, Nivins et al. studied how neonatal non-imaging measures (birth weight [BW], gestational age [GA], and head circumference at birth [HC]) predict imaging-derived measures of brain volume and white matter microstructure at school age (9-10y) (11). Only birthweight showed some predictive power for white matter microstructure via the tract-based spatial statistics (TBSS) analysis in that study. However, regional or white matter skeleton-based approaches often lack the sensitivity and spatial precision of tract based functional analyses, which offer detailed insights into the microstructural characteristics of white matter fiber tracts (12). Building on these results, the goal of the current study was to replicate their findings in a separate sample via an alternative, more sensitive fiber tract-based DTI analysis framework (13).

Here, we aimed to investigate how common neonatal indicators of health (BW, GA and HC) and brain development are related to DTI-based measurement of white matter microstructure in 8 to 10 years old children.

## Materials and Methods

### Participants

The UNC Early Brain Development Study (EBDS) database is a longitudinal dataset that has tracked children from prenatal stages, combining imaging data with cognitive and behavioral assessments throughout the course of their postnatal brain development (14, 15). Parents were recruited from UNC Hospitals and Duke University Medical Center during the second trimester of pregnancy, when they provided written informed consent. Mothers were excluded from the study for major illness or use of illegal drugs during pregnancy. From the EBDS cohort, we retrospectively selected the participants with diffusion MRI scans collected at ages of 8 to 10 years on a 3T Siemens Tim Trio either with a three-shell diffusion MRI sequence (diffusion shell b-value [number of diffusion weighted images per shell]: 0 [13], 300 [8], 700 [32], 2000 [64]; 2x2x2mm^3^ resolution) or a single-shell sequence (b = 0 [7], 1000 [42]), as well as available perinatal clinical information of head circumference, weight, and gestation age at birth. A total of 148 subjects were selected based on the above criteria. All study protocols were approved by the Institutional Review Board of the UNC at Chapel Hill, and written informed consent was obtained from the parents of all participants. Subject recruitment started on 07/09/2003 and ended on 02/13/2008.

### MRI processing and analysis

A study-specific quality control protocol was applied to all raw DTI data using the dmriprep (16) module in the DMRIPlayground toolkit (https://github.com/NIRALUser/DTIPlayground), which includes correction for motion, eddy-current and susceptibility artifacts, as well as rejection of DWI volumes exhibiting significant residual slice-wise or gradient-wise artifacts. After conducting the quality control protocol, 31 subjects were excluded due to insufficient image quality, and limited brain coverage. Brain masks were estimated from the average b=0 image and manually edited. Diffusion tensor images (DTI) were estimated via weighted least-squares. A study-specific DTI atlas was created via the dmriatlas module in DMRIPlayground. Fifty-four major white matter tracts were determined in that DTI atlas space via propagation of the EBDS pediatric DTI atlas (12) followed by automated fiber tracking and tract labeling (17). Profiles of diffusion tensor metrics (fractional anisotropy [FA], axial diffusivity [AD], radial diffusivity [RD]) were extracted at evenly spaced points (arc lengths) along each fiber tract.

Statistical analysis of fiber tract profiles was performed via FADTTS (18), covarying for sex, scan type (single-shell vs three-shell acquisition) yielding tract-wise global p-values as well as local p-value profiles. For each tract, a multivariate analysis was first performed, combining all three DTI metrics in a single model, followed by a posthoc univariate analysis for each FA, RD and AD separately. A summarized overview of the framework can be seen in Figure 1.

**Figure 1.**
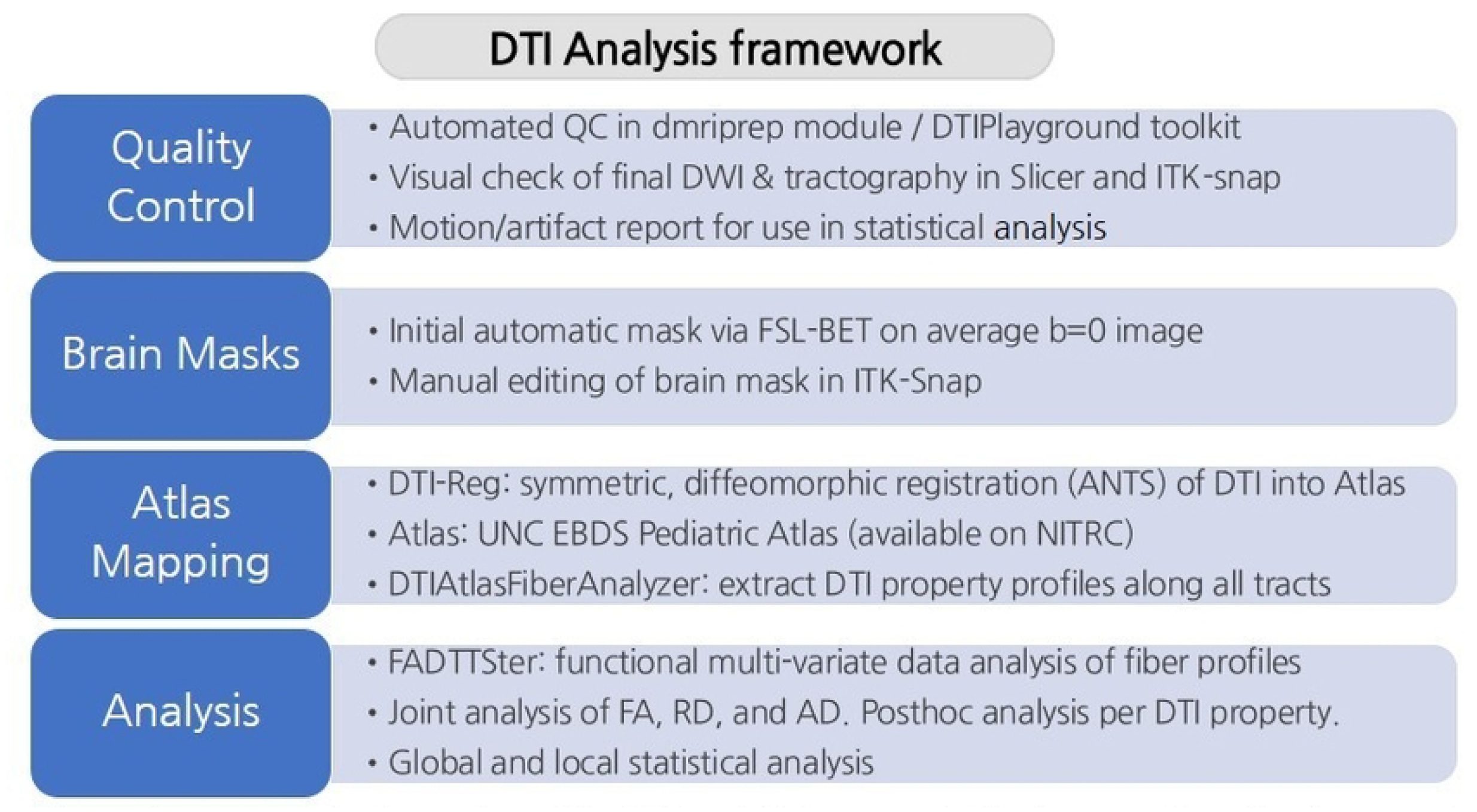
A summarized overview of the DTI analysis framework. The figure outlines the framework from quality control steps to statistical analysis of white matter tracts.

## Results

Out of 148 subjects, 31 were excluded during the quality control protocol, leaving 117 subjects for the final analysis. The baseline characteristics of the 117 subjects included in the final analysis are presented in Table 1. There were 94 subjects using the 3-shell diffusion MRI sequence and 23 subjects using the 1-shell diffusion MRI sequence.

**Table 1.** Demographic characteristics of the children included in DTI analysis. Data for head circumference were available for 113 subjects (4 missing).

| Characteristic | N |  |
| --- | --- | --- |
| Child sex |  |  |
| Male | 57 (48.7%) |  |
| Female | 60 (52.3%) |  |
| Race |  |  |
| White | 93 |  |
| Black or African American | 24 |  |
| Twin status |  |  |
| Non-twin | 52 |  |
| Twin | 65 |  |
| Monozygotic | 38 |  |
| Dizygotic | 27 |  |
|  | Mean<br>( Standard Deviation) | Range |
| Gestational age at birth(weeks) | 36.46 (3.06) | 27.42 - 41.14 |
| Birthweight(g) | 2740.64 (782.03) | 790 - 4732 |
| Gestational age at MRI scan(years) | 9.24 (1.02) | 8.0 - 10.9 |
| Head circumference at birth(cm)* | 32.54 (2.33) | 24.0 - 37.75 |

### Multivariate fiber-tract based analysis

Among the three factors birth weight (BW), gestational age at birth (GA), and head circumference (HC), GA showed the most widespread significant associations (p<0.05) with white matter diffusion metrics globally (38 out of 54 white matter tracts) and locally (see Figure 2 and Table 2). BW (36 out of 54) showed a slightly lower number of globally associated tracts but also showed widespread significant predictive associations. HC (3 out of 54) showed a low number of tracts affected with significant associations. Sex and scan type were also significantly associated with the diffusion metrics globally in some white matter tracts.

**Figure 2.**
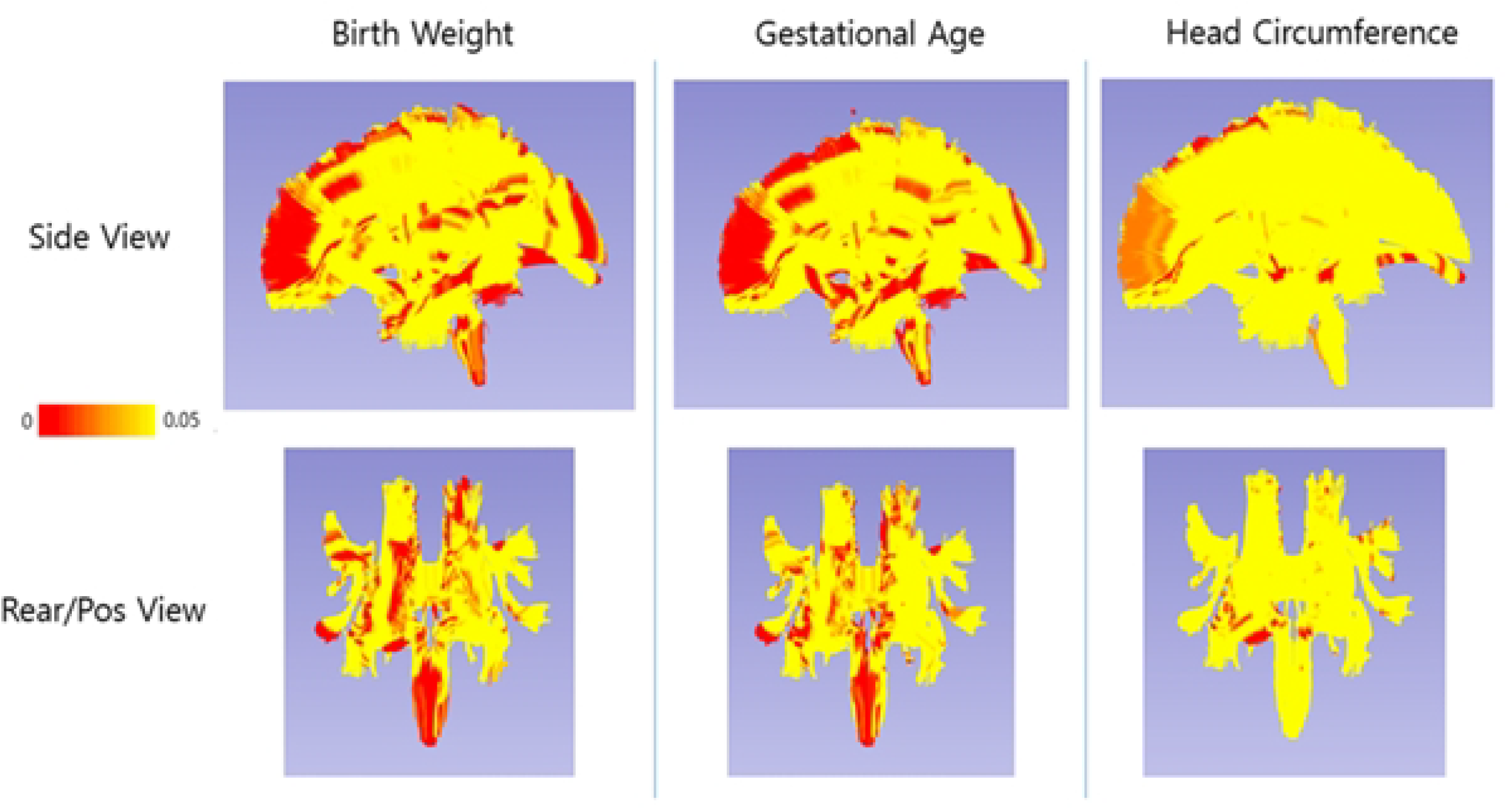
Local fiber tract analysis: Significant correlation (p<0.05) of tensor measures (jointly) with Head circumference at birth, Gestational age and Birthweight for all major 54 tracts. Yellow to red indicates the level of the p values (red = significant).

**Table 2.** Number of tracts showing significant associations (p < 0.05) with perinatal factors. Values are shown as n / total tracts per group. This summary highlights how many tracts within each major white matter pathway were significantly associated with birth weight, gestational age, or head circumference at birth. Full tract-level results are provided in Supplementary Table S1.

| Tract group | Birth Weight | Gestational Age | Head Circumference |
| --- | --- | --- | --- |
| Arcuate Fasciculus | 3/6 | 2/6 | 0/6 |
| Cingulum | 2/4 | 3/4 | 0/4 |
| Corpus Callosum | 5/8 | 5/8 | 0/8 |
| Corticofugal Tracts | 7/8 | 8/8 | 1/8 |
| Corticoreticular Tracts | 2/2 | 2/2 | 0/2 |
| Corticospinal Tracts | 2/2 | 1/2 | 0/2 |
| Corticothalamic Tracts | 7/10 | 7/10 | 1/10 |
| Fornix | 2/2 | 2/2 | 0/2 |
| Inferior Frontoccipital Fasciculus | 2/2 | 2/2 | 0/2 |
| Inferior Longitudinal Fasciculus | 1/2 | 1/2 | 1/2 |
| Optic Radiation | 0/2 | 1/2 | 0/2 |
| Optic Tract | 0/2 | 1/2 | 0/2 |
| Superior Longitudinal Fasciculus | 1/2 | 1/2 | 0/2 |
| Uncinate Fasciculus | 2/2 | 2/2 | 0/2 |
| Total (54) | 36/54 | 38/54 | 3/54 |

### Posthoc univariate analysis

In the posthoc analysis, we found far more correlations for all three perinatal factors with AD than with RD or FA. AD associations with BW, GA and HC were found in 30, 31, 8 out of 54 white matter tracts, respectively. For RD and FA, 5, 13, 3 and 12, 18, 2 out of 54 white matter tracts showed significant associations. For significant tracts, FA and AD were consistently positively associated with BW, GA, and HC, i.e., larger values of BW, GA, or HC lead to higher FA and AD later. RD showed a more variable pattern, without a clear predominance of either positive or negative associations. (see Figure 3 and 4).

**Figure 3.**
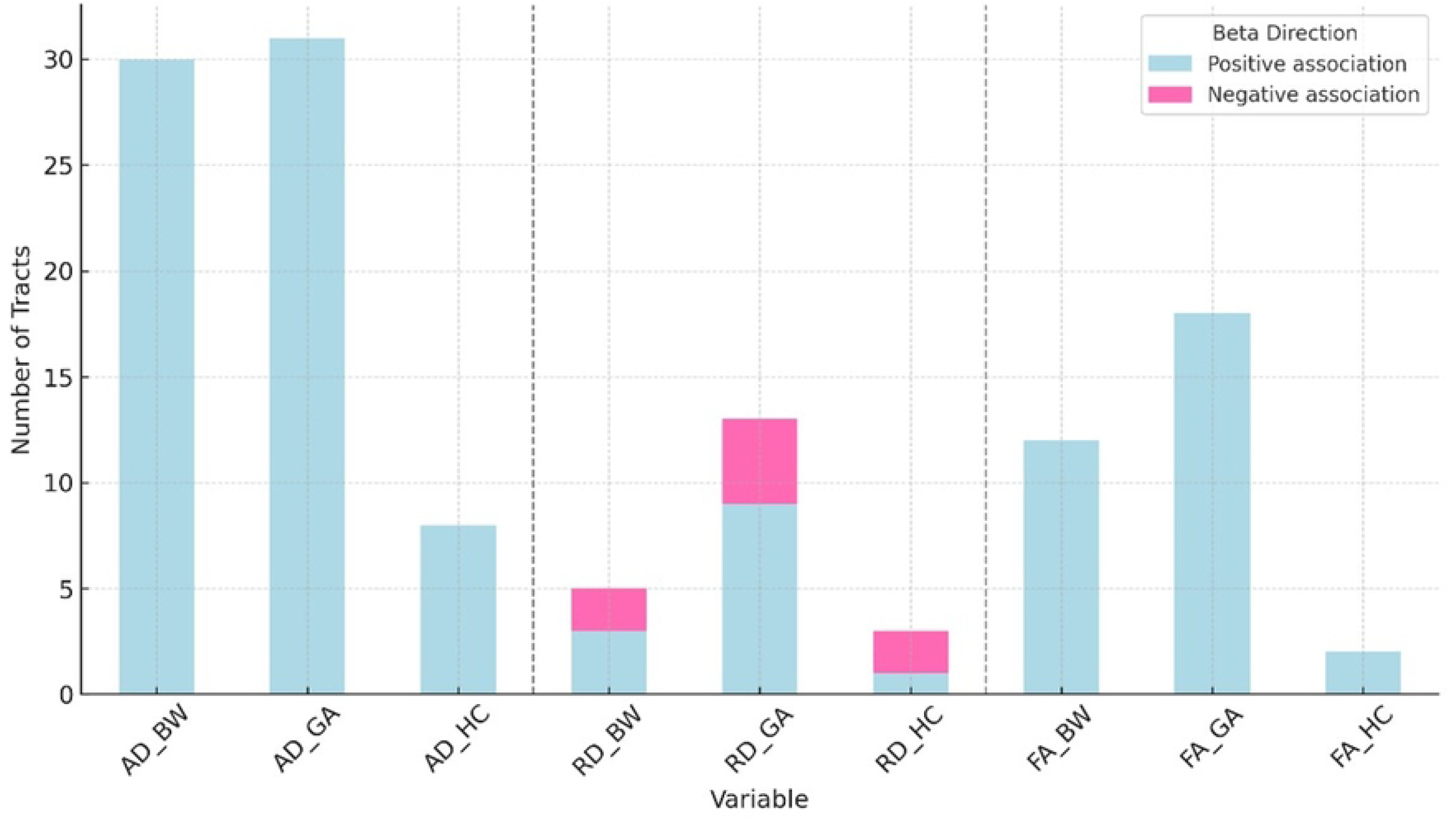
Directionality of beta coefficients for significant tracts associated with BW (birthweight), GA (gestational age), and HC (head circumference at birth), grouped by diffusion metrics (AD, RD, FA). Positive and negative associations are indicated in blue and pink, respectively.

**Figure 4.**
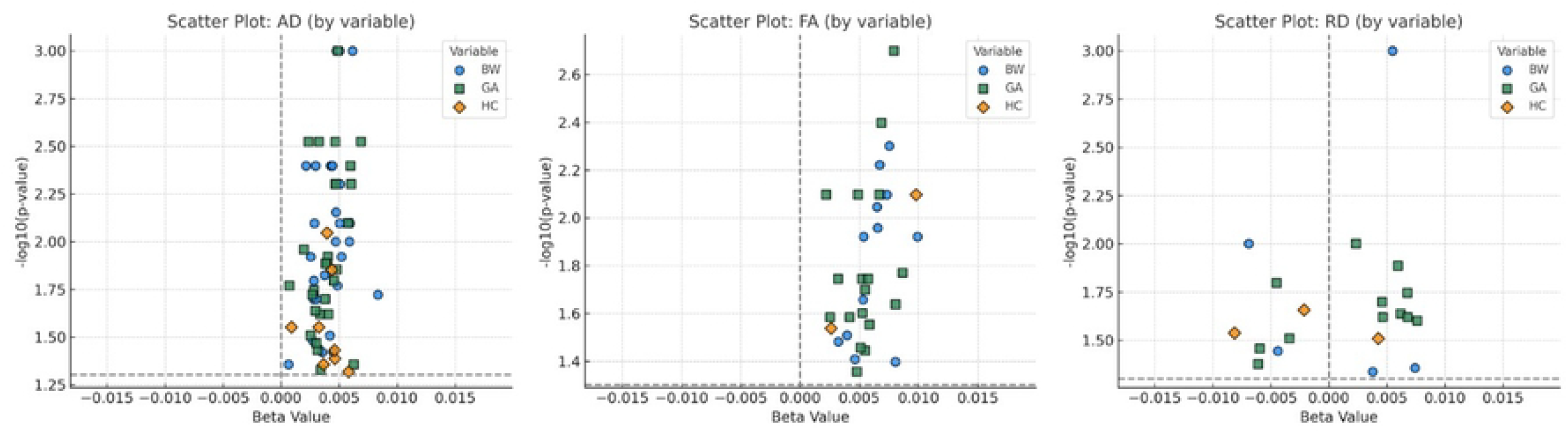
Figure 4 shows scatter plots of beta coefficients vs. –log10(p-value) for each significant tract. Each point represents a significant association, with color and shape indicating the variable (blue circle: BW, green square: GA, orange diamond: HC). The x-axis reflects the direction and magnitude of the association, and the y-axis represents statistical significance.

## Discussion

In this study, we demonstrated that neonatal health indicators, particularly gestational age (GA) and birth weight (BW), are predictive of white matter microstructure at ages 8-10 years as measured by DTI. These associations persisted, albeit weakly, even after accounting for HC. While previous studies have shown associations between perinatal factors and brain volumes in this age group (19-23), our findings extend this understanding to white matter organization.

Our findings both align with and differ from prior research in important ways. While earlier DTI studies using Tract-Based Spatial Statistics (TBSS) found limited associations between perinatal factors and diffusion metrics (11, 24), our fiber-tract analysis framework revealed more widespread relationships. TBSS is well-known to serve well as a hypothesis generation tool due to its whole-brain, white matter skeleton based analysis framework (25, 26), but it is also known to be less sensitive than fiber-tract analysis frameworks such as ours or Automated Fiber Quantification (AFQ) (26-29). This might be one possible factor contributing to a better understanding of the effects of perinatal factors, compared to the previous studies analyzed with TBSS (11, 24).

In contrast to prior studies on structural MRI measures, we observed less predictive power of birth HC with later white matter microstructure. Thus, while HC is a strong predictor of later brain volume measures (22, 23), this is not the case for white matter microstructure. This suggests that while overall head size at birth may influence gross brain volumes, it may have less impact on the local organization of white matter tracts compared to other perinatal factors.

Our sample has a wider range of GA, which might be a factor why our study shows widespread associations for GA in contrast to Nivins et al(11). In our study, including preterm birth was intended to allow for an overall developmental impact across all subjects by also analyzing pre-pubertal outcomes in individuals with low GA.

In the results of the post hoc analysis, axial diffusivity (AD) showed the strongest effects compared to FA and RD. AD can help identify changes in axonal growth, especially during childhood when axonal density and length rapidly increase (30, 31). The sensitivity of AD to axonal integrity makes it valuable for tracking the maturation and organization of white matter throughout early developmental stages. Therefore, we expect that neonatal factors are more predictive of white matter organization than myelination differences indicated by RD (32). Overall, our results suggests that higher GA, higher BW, and larger HC at birth are predictive of advanced white matter organization.

Several limitations warrant consideration. Our moderate sample size (n=117) and use of a single 3T MRI scanner may limit generalizability. Additionally, our cohort’s heterogeneity, spanning preterm to full-term births, while valuable for understanding developmental ranges, introduces potential confounds in developmental trajectories. We also acknowledge that neurodevelopment involves complex interactions between genetic and environmental factors that our analysis cannot fully address. Finally, while the current results accounts for twin status (whether a subject is part of the twin cohort or the singleton cohort), we did not account for the relatedness of when both of the twin subjects were included in the analysis.

Future directions include expanding the study to all participants from the EBDS cohort who have been scanned at ages 8-10, significantly increasing the sample size to over 300 subjects. These scans were conducted using three different 3T Siemens MRI scanners, which will require additional harmonization of the data to ensure consistency across different scanners. In addition, the study plans to implement a more detailed analysis of white matter microstructure using the Neurite Orientation Dispersion and Density Imaging (NODDI) model, which will allow for a more refined understanding of brain development.

In conclusion, we have demonstrated that common perinatal indicators of health (birthweight [BW], gestational age [GA], head circumference at birth [HC]) are associated with white matter organization in 8 to 10 years old children. These findings may provide important insights into our understanding how early life factors impact long-term brain development.

## Acknowledgments

We thank the children and their families who participated in the UNC Early Brain Development Study, and the research staff for their assistance with data collection and management.

S1 Table. Fiber tract-based analysis results showing global p-values for all 54 white matter tracts. Light green shading indicates tracts with significant associations (p < 0.05) with birth weight, gestational age, or head circumference at birth. This table provides the full tract-level results complementing the summary data presented in Table 2.

## References

1. Stiles J. The fundamentals of brain development: Integrating nature and nurture: Harvard University Press; 2008.

2. Mabbott DJ, Noseworthy M, Bouffet E, Laughlin S, Rockel C. White matter growth as a mechanism of cognitive development in children. Neuroimage. 2006;33(3):936–46.

3. Nagy Z, Westerberg H, Klingberg T. Maturation of white matter is associated with the development of cognitive functions during childhood. J Cogn Neurosci. 2004;16(7):1227–33.

4. Filley CM, Fields RD. White matter and cognition: making the connection. J Neurophysiol. 2016;116(5):2093–104.

5. Mukherjee P, Berman JI, Chung SW, Hess CP, Henry RG. Diffusion tensor MR imaging and fiber tractography: theoretic underpinnings. AJNR Am J Neuroradiol. 2008;29(4):632–41.

6. Pecheva D, Kelly C, Kimpton J, Bonthrone A, Batalle D, Zhang H, et al. Recent advances in diffusion neuroimaging: applications in the developing preterm brain. F1000Res. 2018;7.

7. Anjari M, Counsell SJ, Srinivasan L, Allsop JM, Hajnal JV, Rutherford MA, et al. The association of lung disease with cerebral white matter abnormalities in preterm infants. Pediatrics. 2009;124(1):268–76.

8. Ball G, Counsell SJ, Anjari M, Merchant N, Arichi T, Doria V, et al. An optimised tract-based spatial statistics protocol for neonates: applications to prematurity and chronic lung disease. Neuroimage. 2010;53(1):94–102.

9. Chau V, Poskitt KJ, McFadden DE, Bowen-Roberts T, Synnes A, Brant R, et al. Effect of chorioamnionitis on brain development and injury in premature newborns. Ann Neurol. 2009;66(2):155–64.

10. Zwicker JG, Grunau RE, Adams E, Chau V, Brant R, Poskitt KJ, et al. Score for Neonatal Acute Physiology–II and neonatal pain predict corticospinal tract development in premature newborns. Pediatric neurology. 2013;48(2):123-9. e1.

11. Nivins S, Kennedy E, McKinlay C, Thompson B, Harding JE, Children with H, et al. Size at birth predicts later brain volumes. Sci Rep. 2023;13(1):12446.

12. Short SJ, Jang DK, Steiner RJ, Stephens RL, Girault JB, Styner M, et al. Diffusion Tensor Based White Matter Tract Atlases for Pediatric Populations. Front Neurosci. 2022;16:806268.

13. Verde AR, Budin F, Berger JB, Gupta A, Farzinfar M, Kaiser A, et al. UNC-Utah NA-MIC framework for DTI fiber tract analysis. Front Neuroinform. 2014;7:51.

14. Knickmeyer RC, Gouttard S, Kang C, Evans D, Wilber K, Smith JK, et al. A structural MRI study of human brain development from birth to 2 years. J Neurosci. 2008;28(47):12176–82.

15. Stephens RL, Langworthy BW, Short SJ, Girault JB, Styner MA, Gilmore JH. White Matter Development from Birth to 6 Years of Age: A Longitudinal Study. Cereb Cortex. 2020;30(12):6152–68.

16. Dubos J, Park SK, Vlasova R, Prieto JC, Styner M. Dmriprep: open-source diffusion MRI quality control framework with graphical user interface. Proc SPIE Int Soc Opt Eng. 2023;12464.

17. Ngattai Lam PD, Belhomme G, Ferrall J, Patterson B, Styner M, Prieto JC. TRAFIC: Fiber Tract Classification Using Deep Learning. Proc SPIE Int Soc Opt Eng. 2018;10574.

18. Zhu H, Kong L, Li R, Styner M, Gerig G, Lin W, et al. FADTTS: functional analysis of diffusion tensor tract statistics. Neuroimage. 2011;56(3):1412–25.

19. El Marroun H, Zou R, Leeuwenburg MF, Steegers EAP, Reiss IKM, Muetzel RL, et al. Association of Gestational Age at Birth With Brain Morphometry. JAMA Pediatr. 2020;174(12):1149–58.

20. Davis EP, Buss C, Muftuler LT, Head K, Hasso A, Wing DA, et al. Children’s Brain Development Benefits from Longer Gestation. Front Psychol. 2011;2:1.

21. de Kieviet JF, Zoetebier L, van Elburg RM, Vermeulen RJ, Oosterlaan J. Brain development of very preterm and very low-birthweight children in childhood and adolescence: a meta-analysis. Dev Med Child Neurol. 2012;54(4):313–23.

22. Catena A, Martinez-Zaldivar C, Diaz-Piedra C, Torres-Espinola FJ, Brandi P, Perez-Garcia M, et al. On the relationship between head circumference, brain size, prenatal long-chain PUFA/5-methyltetrahydrofolate supplementation and cognitive abilities during childhood. Br J Nutr. 2019;122(s1):S40–S8.

23. Cheong JL, Hunt RW, Anderson PJ, Howard K, Thompson DK, Wang HX, et al. Head growth in preterm infants: correlation with magnetic resonance imaging and neurodevelopmental outcome. Pediatrics. 2008;121(6):e1534–40.

24. Ou X, Glasier CM, Ramakrishnaiah RH, Kanfi A, Rowell AC, Pivik RT, et al. Gestational Age at Birth and Brain White Matter Development in Term-Born Infants and Children. AJNR Am J Neuroradiol. 2017;38(12):2373–9.

25. Smith SM, Jenkinson M, Johansen-Berg H, Rueckert D, Nichols TE, Mackay CE, et al. Tract-based spatial statistics: voxelwise analysis of multi-subject diffusion data. Neuroimage. 2006;31(4):1487–505.

26. Bach M, Laun FB, Leemans A, Tax CM, Biessels GJ, Stieltjes B, et al. Methodological considerations on tract-based spatial statistics (TBSS). Neuroimage. 2014;100:358–69.

27. Yeatman JD, Dougherty RF, Myall NJ, Wandell BA, Feldman HM. Tract profiles of white matter properties: automating fiber-tract quantification. PLoS One. 2012;7(11):e49790.

28. Kuchling J, Backner Y, Oertel FC, Raz N, Bellmann-Strobl J, Ruprecht K, et al. Comparison of probabilistic tractography and tract-based spatial statistics for assessing optic radiation damage in patients with autoimmune inflammatory disorders of the central nervous system. Neuroimage Clin. 2018;19:538–50.

29. Wang X, Zhou C, Wang Y, Wang L. Microstructural changes of white matter fiber tracts induced by insular glioma revealed by tract-based spatial statistics and automatic fiber quantification. Sci Rep. 2022;12(1):2685.

30. Genc S, Raven EP, Drakesmith M, Blakemore SJ, Jones DK. Novel insights into axon diameter and myelin content in late childhood and adolescence. Cereb Cortex. 2023;33(10):6435–48.

31. Dimond D, Rohr CS, Smith RE, Dhollander T, Cho I, Lebel C, et al. Early childhood development of white matter fiber density and morphology. Neuroimage. 2020;210:116552.

32. Song SK, Yoshino J, Le TQ, Lin SJ, Sun SW, Cross AH, et al. Demyelination increases radial diffusivity in corpus callosum of mouse brain. Neuroimage. 2005;26(1):132–40.

